# Microbial Biomarkers of Intestinal Barrier Maturation in Preterm Infants

**DOI:** 10.1101/316257

**Authors:** Bing Ma, Elias McComb, Pawel Gajer, Hongqiu Yang, Mike Humphrys, Adora C. Okogbule-Wonodi, Alessio Fasano, Jacques Ravel, Rose M Viscardi

## Abstract

Intestinal barrier immaturity, or “leaky gut,” is the proximate cause of susceptibility to necrotizing enterocolitis in preterm neonates. However, the impact of intestinal microbiota development on intestinal mucosal barrier maturation has not been evaluated in this population. In this study, we investigated a longitudinally sampled cohort of 38 preterm infants monitored for intestinal permeability (IP) and fecal microbiota during the first two weeks of life. Rapid decrease in IP indicating intestinal barrier function maturation correlated with significant increase in community diversity. In particular, members of the *Clostridiales* and *Bifidobacterium* were highly transcriptionally active, and progressively increasing abundance in *Clostridiales* was significantly associated with decreased gut permeability. Further, neonatal factors previously identified to promote intestinal barrier maturation, including early exclusive breastmilk feeding and low antibiotic exposure, favor the early colonization of the gut microbiota by members of the *Clostridiales*, which altogether are associated with improved intestinal barrier function in preterm infants.

## Introduction

The intestinal mucosa paracellular trafficking of macromolecules is controlled by a competent epithelial barrier ^1^. The intestinal barrier constitutes a protective shield to the diffusion of pathogens and other elements with pro-inflammatory and tissue injury properties, and regulates absorption and secretion of essential nutrients ^2^. A functional intestinal barrier is driven by a complex structure that includes physical barrier with coinciding chemical, immunological and microbiological components ^3^. The colonization with microorganisms starts at birth and undergoes rapid shifts in composition and structure as the host matures over time ^4^. These microorganisms perform essential functions mechanistically linked to intestinal barrier competency, including epithelial metabolism, proliferation and survival, mucin and antimicrobial compound production, and cell-cell communication signaling molecule secretion ^3^. The microbial community in general is considered to play critical roles in the early development of the intestinal epithelium, the immune system, nutrient acquisition and energy regulation, and opportunistic pathogens suppression ^3,5^.

Disrupting intestinal microbiota, on the other hand, leads to dysbiosis, a state of ecological imbalance where the community loses diversity, key bacterial species, and more critically metabolic capacity with reduced colonization resistance to opportunistic pathogens ^6^. Early life gut dysbiosis is associated with disease susceptibility along with short-term and lifelong health issues, such as necrotizing enterocolitis (NEC) ^7^, sepsis ^7^, asthma and allergies ^8^, type 1 diabetes ^9^, celiac disease ^10^, inflammatory bowel disease ^11^ and obesity ^12^, among others. NEC is a life-threatening, gastrointestinal emergency affecting approximately 7-10% of preterm neonates with mortality as high as 30-50% ^13^. In this condition, bacteria across the intestinal wall leading to local and systemic infection and inflammation, and bowel wall necrosis and perforation. Intestinal barrier immaturity, characterized as elevated intestinal permeability (IP), or “leaky gut”, is the proximate cause of susceptibility to NEC in preterm neonates ^14,15^. It is critical to characterize the preterm infant intestinal microbiota to identify dysbiotic states associated with increased intestinal leakiness, as well as beneficial bacteria associated with improved intestinal barrier function, for subsequent stratification of early diagnosis, early intervention and primary prevention of leaky gut and its sequelae.

Despite the critical role of the microbial community in intestinal barrier function, its effect on newborn IP is unknown. In particular, the microbiota of preterm neonates with measured elevated IP, a high-risk population for NEC, has not been studied previously. We hypothesize that the intestinal microbiota plays a pivotal role in modulating IP and that the presence of “beneficial” bacteria will be associated with improved intestinal barrier function in preterm infants. In this study, we studied a cohort of 38 preterm infants born prior to 33 weeks of gestation. IP was measured by urinary detection of orally administered sugar probes lactulose and rhamnose using high pressure liquid chromatography ^16^ with coinciding measures of the composition and function of the fecal microbial communities were investigated. We sampled three time points, study day 1, 8, and 15, during the first two weeks of life, which is a critical period corresponding to the initiation of the intestinal microbiota development process ^16-18^. A rapid decrease in IP was observed to correlate with increased fecal microbiota biodiversity, indicating intestinal barrier function maturation over the first two weeks of life with a shift in the composition and structure in intestinal microbial community. We subsequently revealed an association between decreased IP and the abundance of *Clostridiales*, which was highly transcriptionally active along with members of the *Bifidobacterium*. Our study highlights the multifactorial processes involved in intestinal barrier maturation, and the importance to consider microbiological and neonatal factors for diagnosing, monitoring, and modulating IP in preterm infants.

## Results

### Intestinal barrier maturation correlates with increased microbiota biodiversity over the first two weeks of life

The demographic, obstetric, and neonatal characteristics for all thirty-eight preterm infants enrolled in the study are summarized in **Table 1**. As previously reported ^16^, 20 infants (57%) experienced a rapid decrease in intestinal permeability (IP), 5 infants (14%) had a decreased IP during the first week and subsequent substantial increase and 10 infants (29%) showed a delayed IP decrease maintaining high IP throughout the study period. At each time point, infants were assigned to either high or low IP (**Supplemental File 3**). The microbiota of 64 fecal samples were successfully characterized by high-throughput sequencing of the V3-V4 variable regions of 16S rRNA genes. A total of 422,444 high-quality amplicon sequences was obtained, corresponding to 10,544 (±4,029) sequences per sample with an average length of 428 bp. The top 25 most abundant phylotypes are shown in **Supplemental Figure 1A**. Taxonomic profiles of all samples were clustered into three distinct groups according to similarities in community composition and structure. *Klebsiella spp.*, *Staphylococcus epidermidis*, and *E. coli* dominated cluster I, II, and III, respectively. Both La/Rh ratio and taxonomic profile of each sample are shown in **Supplemental File 3**. Taxonomic profiling of corresponding metagenomes further resolved *Klebsiella* spp. to *Klebsiella pneumoniae*. Not surprisingly, older term infants at 6-24 months old, or phase II/III as defined previously ^17-19^, clustered together in a different and more diverse cluster (**Supplemental Figure 1B**). Rapid decrease in IP over the two-week observation period indicates intestinal barrier function maturation (p-value = 0.002), which is correlated with a significant increase in community diversity (p-value = 0.02) (**Figure 1A**); while delayed increase in community diversity was associated with maintenance of high intestinal permeability (p-value < 0.001) (**Figure 1B**). The results indicated that preterm infant intestinal barrier maturation correlates with increased fecal microbiota biodiversity and a change in microbiota structure.

**Figure 1.**
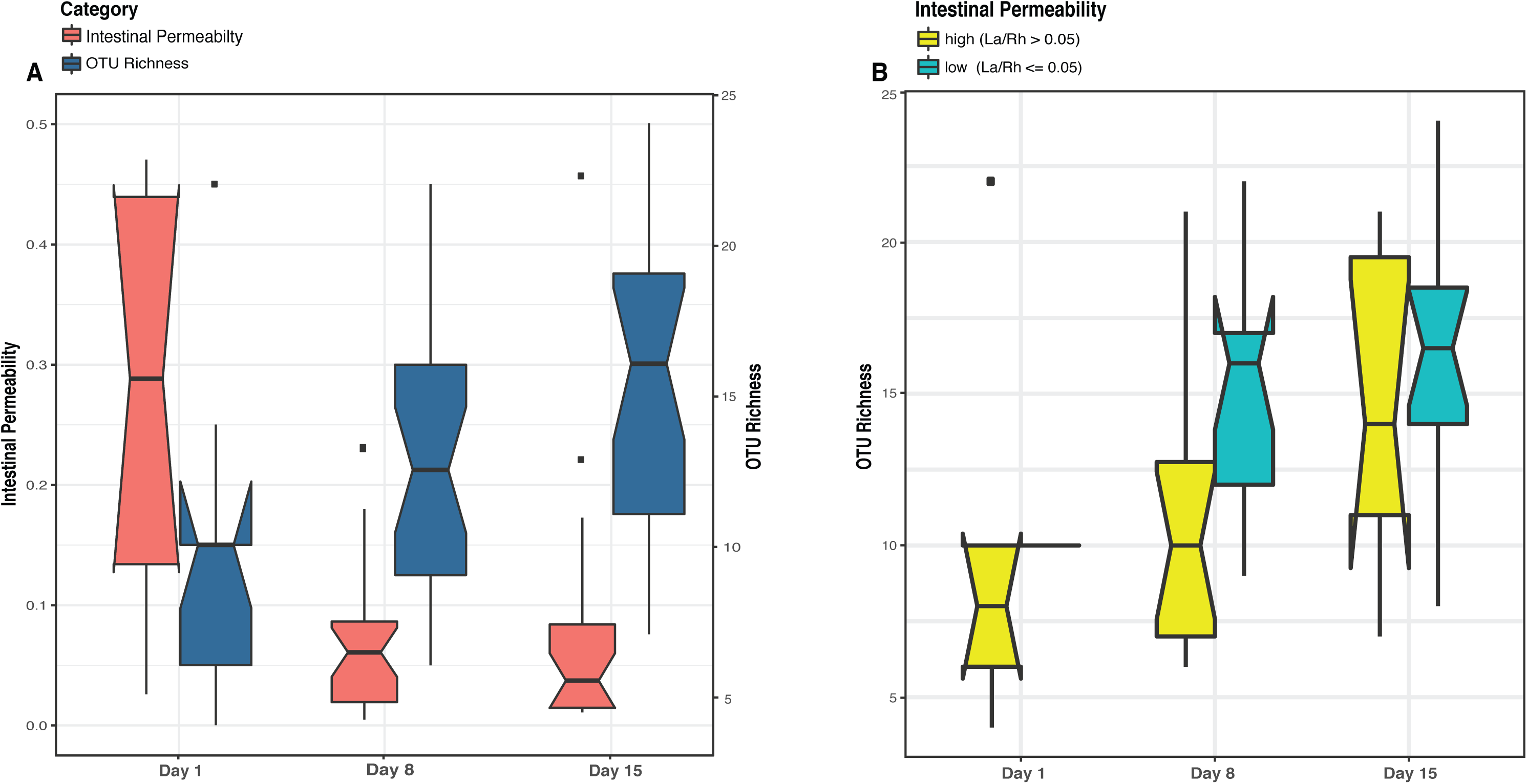
Boxplots comparing levels of intestinal permeability and microbial community diversity at study days 1, 8, and 15 in a cohort of 43 preterm infants (<33 weeks gestational age). Intestinal permeability is measured by non-metabolized sugar probes lactulose (La) (marker of intestinal paracellular transport)/rhamnose (Rh) (marker of intestinal transcellular transport). Microbial community diversity was calculated by OTU (Operational Taxonomic Units) richness. Wilcoxon rank sum test and a false discovery rate of 5% was used in significance test. Median values and interquartile of the values were shown in box. (**A**) Intestinal permeability (p-value = 0.002) and community diversity at the three study time points (p-value = 0.02). (**B**) Community diversity (p-value < 0.001) in infants with low and high intestinal permeability defined by a La/Rh >0.05 or <=0.05 respectively (1).

**Table 1.** Characteristics of Study Subjects (preterm infants <33 weeks gestational age) (n=38).

|  | N | % |
| --- | --- | --- |
| **Ethnicity** |  |  |
| African American | 22 | 57.9 |
| Asian | 3 | 7.9 |
| White | 12 | 31.6 |
| Others | 1 | 2.6 |
| **Gender** |  |  |
| Female | 17 | 44.7 |
| Male | 21 | 55.3 |
| **Delivery route** |  |  |
| Cesarean | 26 | 68.4 |
| Vaginal | 12 | 31.6 |
| **Gestational age (Mean, Median, SD)** |  | **(29.9, 31, 2.2)** |
| <= 28 weeks | 10 | 26.3 |
| > 28 weeks | 28 | 73.7 |
| **Birth weight (Mean, Median, SD)** |  | **(1386, 1472, 404)** |
| < 1500 grams | 21 | 55.3 |
| >= 1500 grams | 17 | 44.7 |
| **Antibiotic use** |  |  |
| None | 7 | 18.4 |
| 1 to 3 days | 12 | 31.6 |
| > 4 days | 19 | 50.0 |
| **Day start breastmilk feeding** |  |  |
| Day 1 | 17 | 44.7 |
| Day 2 or 3 | 15 | 39.5 |
| > Day 4 | 6 | 15.8 |
| **Day reached full breastmilk feeding** |  |  |
| < Day 7 | 5 | 13.2 |
| Day 8 to 14 | 15 | 39.5 |
| > Day 15 | 18 | 47.4 |
| **Intestinal permeability pattern** |  |  |
| normal | 24 | 63.2 |
| late increase | 6 | 15.8 |
| delayed | 4 | 10.5 |

### Subject variation, PMA and IP explain most of the variation in microbial community composition

We employed multivariate response linear regression on the “balance” of microbial community and evaluated the effect of covariates of demographic, obstetric, and neonatal factors on the microbiome using Gneiss ^20^. Covariates of antibiotics use, maternal antibiotics use, delivery mode, PPROM, feeding pattern, IP, birthweight, gender, ethnicity, gestational age (GA) and postmenstrual age (PMA) were included in the analysis. Subject, PMA, and IP had the greatest correlation with intestinal microbiota, together they explained 63.4% of the variation of the intestinal microbial community composition observed in the cohort (**Supplemental File 4**). The plots (**Supplemental Figure 2**) show the predicted points lie within the same region as the original communities and the residuals have roughly the same variance as the predictions within ±2. Overall our result indicates the microbial differences between subjects are large (R squared difference is 0.23±0.10), and the covariate with strongest effect is PMA (R squared difference is 0.44). IP correlates with the intestinal microbiota (R squared difference is 0.20), and its effect is lower than PMA and similar to the average among-subject difference.

### *Clostridiales* is associated with low intestinal permeability in preterm infants

Comparative analysis of fecal microbiota with high and low IP showed that *Clostridia*, the class containing the order *Clostridiales* in this cohort, was significantly more abundant in samples with low IP compared to those with high IP (p-value = 0.01) (**Figure 2**, **Supplemental Figure 3**). In particular, a progressive and significant increase in members of *Clostridiales* over the first two week after birth significantly associated with low IP (p-value = 0.0002) (**Figure 3**). Based on Bayesian nonparametric adaptive smoothing models and subject-specific changes in relative abundance of *Clostridiales* at each of study day 1, 8, and 15, the results demonstrated: (1) at baseline study day 1, the abundance of *Clostridiales* was low in subjects with either high or low IP; (2) However, in samples measures with low IP but not high IP, a significant increase in *Clostridiales* was observed that reached ~8% median and >20% maximal relative abundance from study day 1 to day 8, and ~16% median and ~45% maximal relative abundance again from study day 8 to 15; (3) on the other hand, in samples measured in high IP at study day 1 that are also high at the follow up days, members of *Clostridiales* was almost completely absent on study day 8 and no increase was observed from study day 1 to 8, and the increase from study 8 to 15 was small at ~3% median and ~10% maximal relative abundance; (4) in infants 6-24 months old, *Clostridiales* is the most abundant taxonomic groups with >50% median and >85% maximal relative abundance (**Supplemental Figure 4**). Together, our results suggest preterm infants at birth have low abundance of *Clostridiales*, which became progressively and significantly more abundant only in the group with rapid progression of intestinal barrier maturation, while remained low in those with persistent high IP over the first two weeks of life.

**Figure 2.**
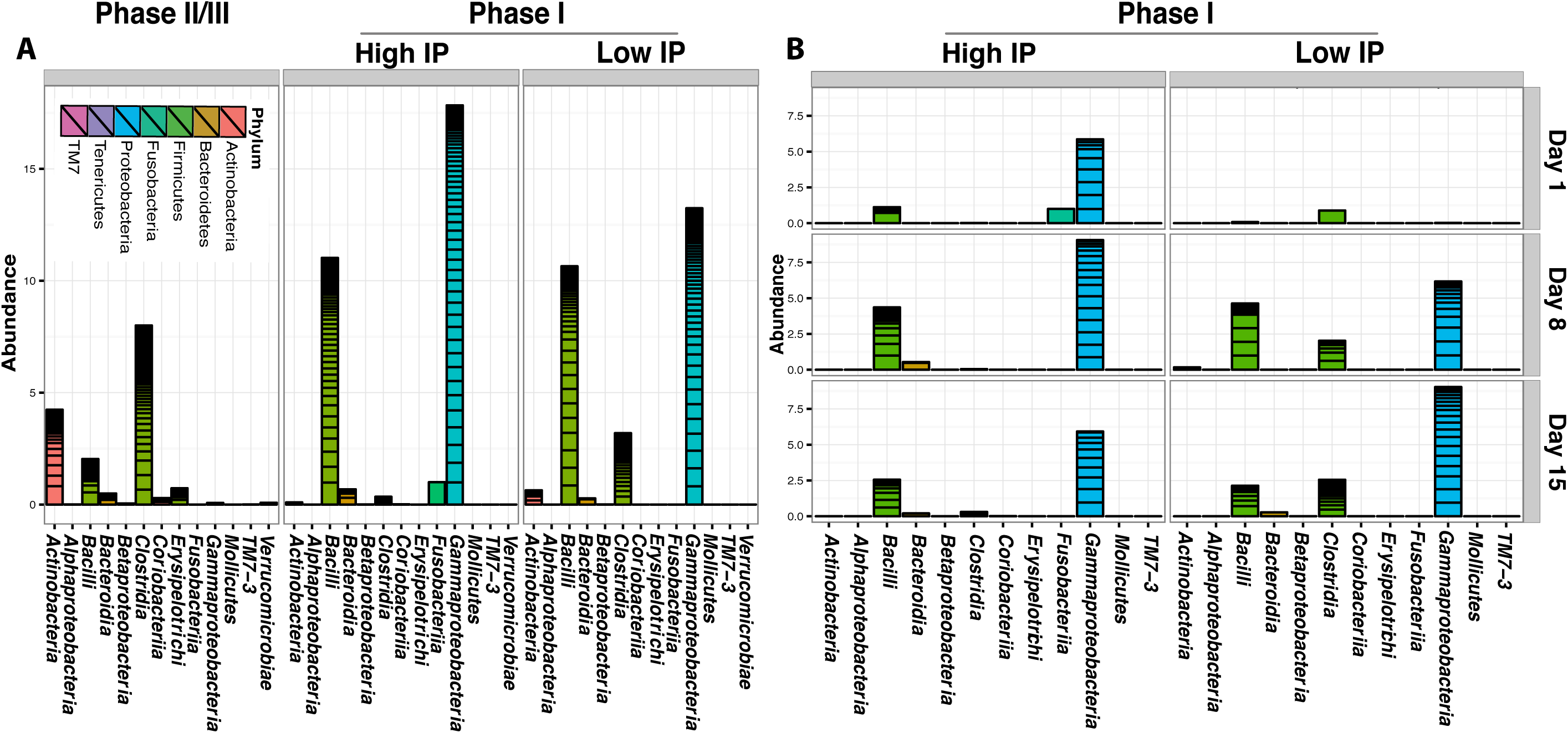
Cumulative relative abundance of bacterial groups in high and low IP infants. (**A**) cumulative abundance between phase II/III subjects (6-24 months of age) and phase I infants (within first two weeks of life) with high and low IP; (**B**) Cumulative abundance at different study day at day 1, 8, and 15 for phase I infants with high and low IP. The most outstanding difference between high and low IP in preterm infants is in the *Clostridiales* (p-value = 0.01), which is the most abundant bacterial group in phase II/III infants.

**Figure 3.**
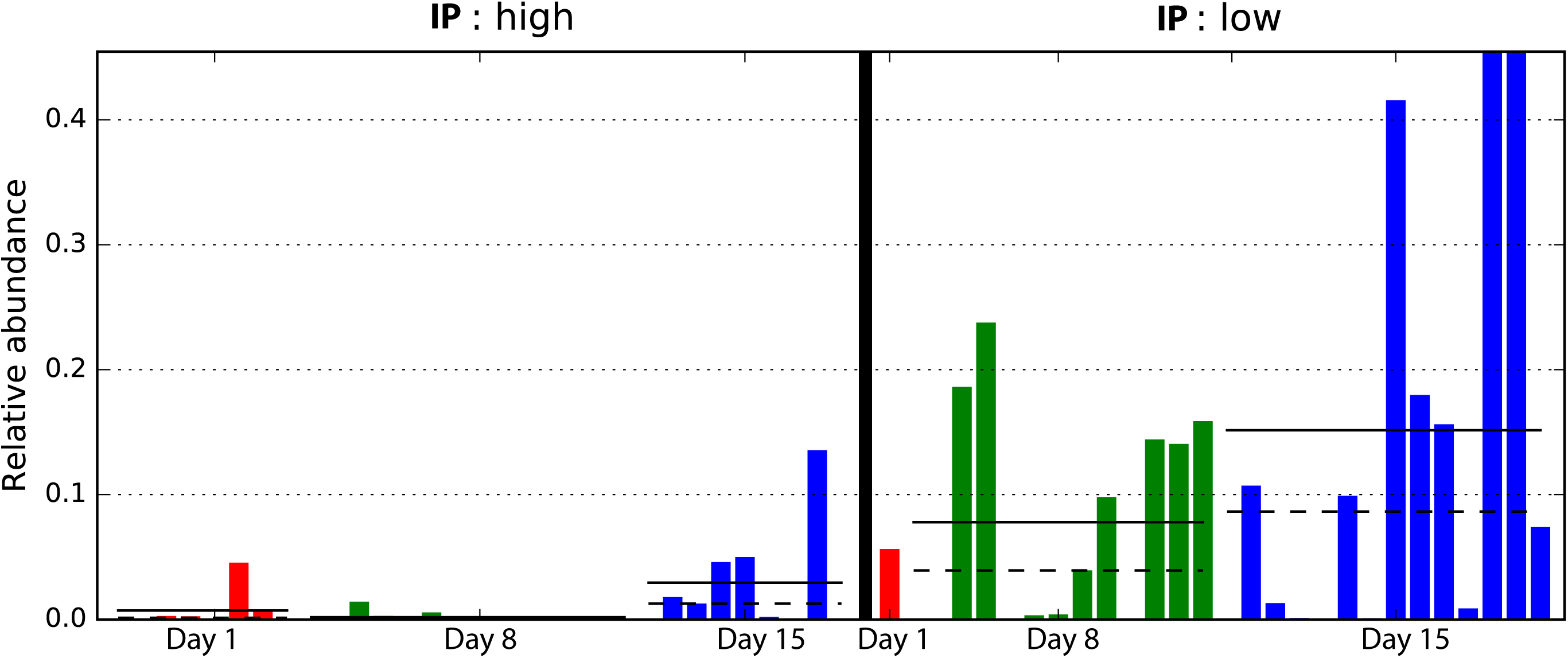
Comparison of samples with different intestinal permeability on the relative abundance of members of *Clostridiales*. Bars represent the relative abundance of *Clostridiales* in each sample. Dotted line represents mean, solid line represents median relative abundance. The alpha value for the non-parametric factorial Kruskal-Wallis sum-rank test was 0.05 and the threshold for the logarithmic LDA model (3) score for discriminative features was set at 2.0. Low IP: La/Rh < 0.05; high IP: La/Rh >= 0.05.

We further calculated the predictive power of microbiota composition in classifying IP using random forest supervised machine learning scheme. The top 15 phylotypes with the highest mean decrease gini index importance measure (**Supplemental Figure 5**) were used to fit a random effect logistic regression model of IP, 4 of which resulted significantly associated with low IP (**Supplemental File 5**), including three members of the order *Clostridiales*, *Coprococcus* (p-value = 0.004), *Lachnospiraceae* (p-value = 0.007), *Veillonella dispar* (p-value = 0.01), and *Bifidobacterium* (p-value = 0.01) from the order of *Bifidobacteriales*. Interestingly, *Bifidobacteriales* was the second most abundant taxonomic groups in infants 6-24 months old, only lower than *Clostridiales* (**Supplemental Figure 4**).

### *Clostridiales* and *Bifidobacterium* are highly active members of the intestinal microbiome

The level of bacterial transcriptional activities was characterized by studying the suite of genes present and expressed in preterm infant intestinal microbiota. A total of 869 million metagenomic sequence reads (average of ~31.0 million sequence reads per sample) and 694 million metatranscriptomic sequence reads (average of ~53.4 million sequence reads per sample) were obtained after quality assessment. **Figure 4** shows that *Bifidobacterium breve* (*Actinobacteria*), *Veillonella* and *Clostridiales Family XI incerteae Sedis* (*Clostridiales*) are the most transcriptionally active bacteria with high ratio of transcript abundances over gene abundances in all samples. Further, the levels of transcriptional activities of *Bifidobacterium breve* and *Clostridiales Family XI incerteae Sedis* are correlated with a spearman correlation of 0.89, suggesting these two taxonomic groups are either functionally dependent or co-regulated (**Supplemental Figure 6)**. We observed increased abundance of both *Clostridiales* and *Bifidobacteriales* through the transition from the first two weeks (phase I) to later age of 6-24 months (phase II/III) as further supporting their active contribution to the function of the GI microbiota after birth. Interestingly, *Clostridiales* and *Bifidobacteriales* are also the most abundant taxonomic groups in the intestinal microbiota of 6-24 months old infants (**Supplemental Figure 4**, **Supplemental File 3**). Specifically, members of the family *Clostridiales* have an average abundance of 50±3% in phase II and III infants, compared to 0.1±0.4% in phase I infants. *Bifidobacteriales* have an abundance of 26±5% in phase II and III as opposed to 0.1±0.3% in phase I infants. Together with the previous observation that *Coprococcus* (*Clostridiales*), *Lachnospiraceae* (*Clostridiales*), *Veillonella dispar* (*Clostridiales*), and *Bifidobacterium* (*Bifidobacteriales*) are significantly associated with low IP, our results suggest the presence and more importantly the activity of bacterial members of *Clostridiales* and *Bifidobacteriales* are associated with improved intestinal barrier function.

**Figure 4.**
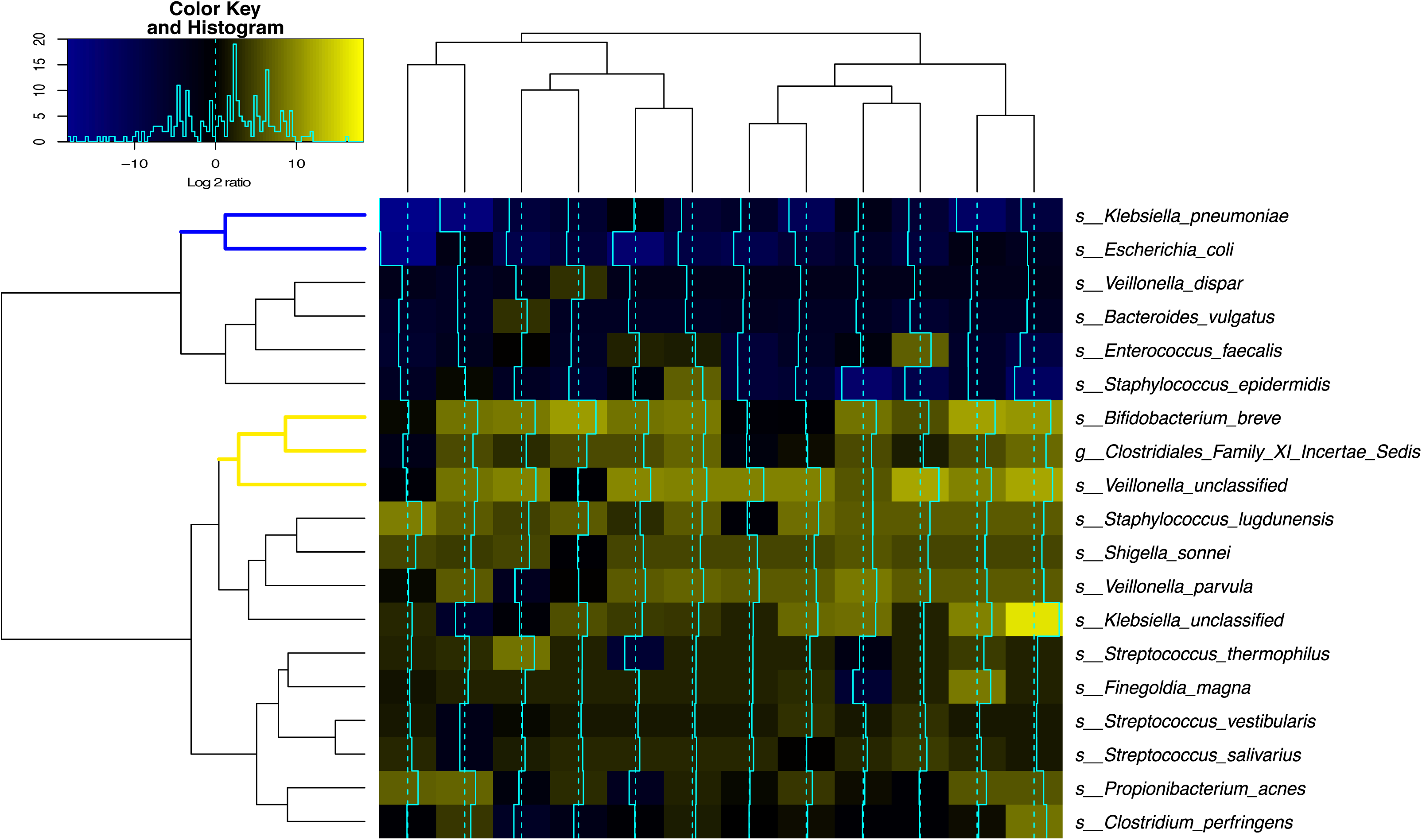
Bacterial species transcriptional activity in preterm infant stools. Fecal samples are represented in columns and taxonomic composition quantified using MetaPhlAn (55) version 2 are shown in rows, both are organized by hierarchical clustering. Normalization using Witten-Bell smoothing was performed, and the relative expression of a gene in a sample was calculated by normalizing the smoothed value of the expression level in the metatranscriptome by the smoothed value of the corresponding gene abundance in the metagenome (56, 57). Color scheme indicates an approximate measure of the species’ clade-specific transcriptional activity (56). The colored branches show the clustering of bacterial species that are consistently transcriptionally active (yellow) or consistently transcriptionally inactive (blue) across samples.

Conversely, the two *Enterobacteriaceae* species, *Klebsiella pneumoniae* and *Escherichia coli*, had low transcriptional activity despite their high relative abundance in the infant GI microbiota, questioning their functional contribution to the infant stool microbiota. Interestingly, *Enterobacteriaceae* and *Staphylococcus* are the most abundant bacterial taxa present in phase I infants but are rarely observed in phase II and III, suggesting their presence in the infant stools is temporary and might not contribute greatly to the functions provided by the GI microbiota.

### Early breast milk feeding and low antibiotic exposure positively correlates with Clostridiales abundance and activities

The associations between intestinal microbiota and demographic, obstetric, and neonatal factors were also evaluated. Gneiss analysis suggests delivery mode, PPROM, gender, ethnicity, birthweight, maternal antibiotics use are not contributing covariates to the intestinal microbial community variance. Further, no bacterial phylotype was identified to significantly associate with these factors. However, breast milk feeding pattern and antibiotic exposure were significantly associated with increased abundance of *Clostridiales.* More specifically, early full exclusive breast milk feeding by study day 10 (p-value = 0.0001) (**Supplemental Figure 7**) and antibiotic exposure limited to no more than 4 days (p-value = 0.05), were associated with the family *Lachnospiraceae* in the *Clostridiales* (p-value = 0.004) (**Supplemental Figure 8**). On the other hand, *Enterobacteriaceae*, particularly *Klebsiella pneumoniae* (as identified by metagenomics sequencing), was significantly associated with full breast milk feeding achieved after study day 10 (p-value = 0.01) (**Supplemental Figure 7)**. These results strongly suggest members of the *Clostridiales* are significantly associated with low intestinal permeability, early full breast milk feeding, as well as shorter duration of antibiotic use.

We further characterized the genotypic variation of *E. coli* through reconstructing MLST loci-sequences from metagenomes ^21^, and compared them to 25 recently characterized *E. coli* MLST genotypes associated with NEC ^22,23^. Five *E. coli* genotypes were only observed in samples with high IP, two of which, sequence type 73 and 131, were previously identified as uropathogenic *E. coli* (UPEC) strains associated with NEC and infant mortality ^23^ (**Supplemental File 6**). One *E. coli* genotype 697 which was not recognized as a UPEC *E. coli* strain nor was observed in NEC ^23^, was observed in both high and low IP samples. Two new MLST genotypes of *E. coli* were also observed. A minimum spanning tree on the sequence types is shown in **Supplemental Figure 9** and was used to demonstrate the relationship among genotypes of *E. coli*.

### *Clostridiales* are highly prevalent in the GI microbiota of preterm infants

The most abundant bacterial species included *K. pneumoniae*, *Staphylococcus epidermidis*, *E. coli*, and *Enteroccocus faecalis* were found with mean abundance of ~10-35% (S.D. ~15%-30%) and ~85-95% prevalence in these samples. In comparison, many species such as *Streptococcus agalactiae*, *B. breve*, *B. longum*, *Clostridium perfringens*, *Propionibacterium acnes, Bacteroides fragilis*, *Veillonella parvula*, and *Streptococcus thermophiles* were present in 15-70% of all samples and had a much lower level of abundance ranging from ~0.0001% to 1% (S.D. ~0.0001%-6%). Many of the members of *Clostridiales* were not resolved at the species or genus-level, while those taxonomically identified *Clostridiales* included *Coprococcus*, *Blautia*, *SMB53*, *Ruminococcus gnavus*, *Clostridium* spp*., Faecalibacterium prausnitzii*, *Dorea*, *Ruminococcus bromii*, *Roseburia*, *Pseudoramibacter* and *Butyricicoccus pullicaecorum* were detected in low or extremely low abundance yet high prevalence (**Supplemental File 3**). A previous study revealed that stool bacterial load varies greatly in the first few days of life but then reached and persisted in most infants in the range of 10^9^ to 10^10^ bacteria per gram of stool after one week of life ^24^. Given our average sequencing depth is ~10^4^-10^5^ it is likely that some bacterial taxonomic groups with low relative abundance (<0.001%) are below our detection limit, and their prevalence is underestimated. It is expected that the prevalence of the members of *Clostridiales* can be higher than the currently observed 15-70% among samples in the GI microbiota of preterm infants. The marked discrepancy between bacterial abundance and prevalence suggests that bacterial species present in the intestinal microbiome of preterm infants can selectively colonize and grow under nutritional or antibiotic pressures.

## Discussion

Preterm infants are at elevated risk for leaky gut, feeding intolerance, NEC and sepsis, and other short-term and long-terms morbidities ^19^. The pathophysiology of these disorders is likely multifactorial, involving a combination of intestinal mucosa barrier immaturity, imbalance in microvascular tone, aberrant microbial colonization and altered immune responses ^19,25,26^. Previously, our group and others demonstrated that neonatal factors such as gestational age, antibiotic exposure, and exclusive breastmilk feeding affect intestinal mucosa barrier permeability in preterm infants ^16,27^. With the rapid development of high-throughput sequencing technology, recent studies have evaluated the significant association between the composition of intestinal microbiota, neonatal intestinal health and development ^3,5,24^. However, the relationships between intestinal microbiota and IP have not yet been evaluated in a high-risk preterm population. In this study, we investigated the early development of the intestinal microbiota and its association with IP in a cohort of 38 preterm infants sampled during the first two weeks of life. We observed that neonatal factors known to be associated with low IP, including early exclusive breast milk feeding and low antibiotic exposure, favored the early colonization of the gut microbiota by members of *Clostridiales*. The associations between neonatal factors, intestinal microbiota and intestinal barrier function further substantiate the multifactorial processes involved in gut barrier maturation, thus highlighting the impact of neonatal care practices and the potential for therapies such as rationally designed live biotherapeutics strategies to rapidly lower IP after birth in preterm infants.

A critical value of understanding the driver of IP, including associated microbiological biomarkers, is in its clinical significance in NEC risk diagnostics and disease prevention. The etiology of NEC involves the interaction between immature intestinal barrier and the developing intestinal microbial community that leads to an excessive inflammatory response ^25,26,28,29^. Though IP is high at birth in preterm infants, it rapidly decreases over the first few days, which is associated with diminished risks for adverse outcomes ^16,30^. Aberrant intestinal barrier function manifests by persistently high and/or late decrease in IP and is likely due to the physiological immaturity of the GI tract barrier function and altered levels of the normal microbial communities ^14,15^, resulting in microbial invasion of the intestinal wall and gut lamina propria triggering a cascading inflammatory response and ultimately intestinal necrosis and severe infection ^2^. Multiple studies have revealed microbial community dysbiosis is involved in stimulating a hyperinflammatory response that leads to NEC ^25,26,28,29^. This community dysbiosis has been characterized by the presence of members of the family *Enterobacteriaceae* such as *E. coli*, *K. pneumonia,* as well as *Enterobacter cloacae* ^23,26,31^. However, a generalized bacterial dysbiosis alone does not adequately explain NEC. Many preterm infants that are colonized by high abundance of *Enterobacteriaceae* do not develop NEC ^32^, and many NEC cases lacked intestinal colonization of *Enterobacteriaceae* ^33^. In this study, *Enterobacteriaceae* was significantly associated with both elevated IP and later attainment of full exclusive breastmilk feeding (>10 days), while other beneficial bacteria such as members of *Clostridiales* and *Bifidobacterium* were significantly associated with improved IP and earlier breastmilk feeding attainment. These results emphasize the importance of a holistic understanding of the etiology of NEC, including the mechanistic characterization of the functional synergy and/or competition among different bacterial groups, as well as nutritional factors, drug uses and host genetics. Further, the links established by previous microbiota association studies could not elucidate the causalities between gut microbiota and NEC development. Our study prospectively associates maturation of gut barrier function with specific microbial community composition and structure for the first time, prior to the onset of NEC. Research on neonatal IP will not only further our understanding of NEC etiology but will help identify the “window of opportunity” for intervention prior to the onset of NEC. Early prediction and prevention of NEC will ultimately improve overall infant survival rates.

Multiple intrinsic and extrinsic factors affect newborn intestinal microbiota, such as maternal diet, delivery mode, breast milk feeding, antibiotic exposure, and other early life environmental exposures ^7,34^. In this study, early exclusive breast milk feeding and low antibiotic exposure was associated with the presence of members of *Clostridiales* in the stool microbiota of preterm infants. We have previously observed these two factors are associated with improved IP in preterm infants ^16^, which has been shown to be critically protective against NEC ^35^. This observation emphasizes the importance of factors such as clinical administration of nutritional supplement and limiting exposure to antibiotic in neonatal care units. Interestingly, *Clostridiales* strains were recently shown to be sensitive to many antibiotics, including ampicillin and amoxicillin ^36^, both commonly used for the neonate clinical management. Further understanding of the selective nutritional requirement that favor the growth of these bacteria would afford the development of novel nutritional supplemental strategies to limit the incidence of NEC and improve clinical outcomes in preterm infants.

Current therapies for NEC are mostly ineffective, and involve antibiotic treatment and surgical interventions, including drain placement or bowel resection. These procedures are associated with poor prognosis and a mortality rate of ~50% due to the rapid progression of the disease ^37^. Live biotherapeutics products (LBP) are being considered, but selecting the appropriate one remains a major challenge. LBP therapies are promising, low-cost, and constitute a likely safe preventive measure to improve intestinal barrier maturation and reduce NEC incidence in at-risk preterm infants ^38^. In an experimental mouse model of NEC, the administration of *Bifidobacterium infantis* prevented an increase in IP, stabilized tight junction proteins, and reduced NEC incidence ^28^. Translating these findings in human has been challenging. There have been at least 11 randomized controlled trials and a recent meta-analysis of LBP supplementation to prevent NEC in preterm neonates ^39^. Although there was a 30% reduction in NEC incidence in these trials, various formulations, doses, and duration of therapy were used, infants <1000 g BW with the highest NEC incidence were under-represented, and no Food and Drug Administration-approved products are available to assure quality and safety under good manufacturing practices.

*Clostridiales* offer a new opportunity to develop a LBP for the prevention of NEC, in combination with strains of *Lactobacillus* and *Bifidobacterium* already available. Members of the family *Clostridiales* often have anti-inflammatory properties associated with their fermentative metabolism of carbohydrates and amino acids ^40^. Because of the difficulties to culture *Clostridiales*, it has been largely overlooked. A few species belonging to this family are known for their pathogenicity and include *C. botulinum*, *C. perfringens*, *C. tetani*, and *C. difficile* ^41^, however these are opportunistic pathogens and not commensal of the intestinal microbiota. The application of culture-independent high-throughput sequencing identified many formerly unculturable *Clostridiales* species, and the group is now thought to be one of the predominant groups of microbes inhabiting the GI tract, comprising ~30-40% abundance of the adult intestinal microbiota ^42^. These species form the basis of the microbiome therapeutics product, SER109, for the treatment of *C. difficile* infection in adults ^43^. *Clostridiales* are heterogeneous in terms of their enzymatic, and metabolic properties, and produce beneficial short-chain fatty acid (SCFA) such as acetate, propionate, and butyrate ^44^. Further, *Clostridiales* have been shown to stimulate the production of intestinal epithelial cytokines that have been associated with the improvement of intestinal dysbiosis, and marked reduction in inflammation ^36,45,46^. The recent characterization of 46 strains of newly isolated *Clostridiales* revealed their ability to induce regulatory T cells and a protection against colitis and allergic responses ^45^. Seventeen strains of human-derived *Clostridiales* species were rationally selected using gnotobiotic mice and the cocktail shown to have prophylactic effect in mouse colitis ^36,46^. In addition, the administration of *Clostridiales* protects the host from pathogen infection and abrogated intestinal pathology ^47^. In term infants, the presence of *Clostridiales* in the intestinal microbiota was demonstrated to prevent colonization by bacterial pathogens such as *S. Typhimurium* ^48^. Unfortunately, the current standard application of 16S rRNA V4 or V3-V4 amplicon sequencing is not capable to resolve the species of *Clostridiales* present in a sample ^49^. Future taxonomic and functional characterization of *Clostridiales* species will greatly improve our capability to develop novel diagnostic and treatment strategies, and potentially prevent microbial community-mediated intestinal dysbiosis in preterm infants to optimize intestinal maturation and limit the burden of prematurity ^23^.

## Methods and Materials

### Participants and intestinal permeability measurement

The institutional review boards of the University of Maryland and Mercy Medical Center approved the study protocol and informed consent was obtained from parents for participation of their infants in the study. All methods were performed in accordance with the relevant guidelines and regulations. Thirty-eight preterm infants 24^0/7^-32^6/7^ weeks GA were enrolled within 4 days after birth and received 1 ml/kg of the non-metabolized sugar probes lactulose (La) (marker of intestinal paracellular transport)/rhamnose (Rh) (marker of intestinal transcellular transport) (8.6 g La +140 mg Rh/100 mL) enterally on study days 1, 8 ± 2 and 15 ± 2. La/Rh was measured by high-pressure liquid chromatography (HPLC) in urine collected over a 4h period following administration of the sugar probes as previously described ^16^. High or low intestinal permeability was defined by a La/Rh >0.05 or <=0.05 respectively, as validated and applied previously ^16^. PMA was calculated as gestational age at birth plus week of life as defined previously ^50^. Fecal samples (~1g) were collected at the same time, stored immediately in 2 ml of RNAlater (QIAGEN). Urine and fecal samples were archived at −80°C until processed. A standard feeding protocol was used for all preterm participants. To compare microbiota of infants at different growth phases ^17,19^, 16 samples from older term infants at phase II/III (6-24 months old) from a previous study ^51^ were included in the comparative analyses.

### Stool nucleic acid extraction and sequencing

DNA was extracted from all samples as previously reported ^52^. Briefly, a 500 μl aliquot of fecal material mixture was homogenized and carefully washed twice in PBS buffer. Enzymatic lysis using mutanolysin, lysostaphin and lysozyme was performed, followed by proteinase K, SDS treatment and bead beating. DNA purification from lysates was done on a QIAsymphony automated platform. PCR amplification of the V3-V4 variable region of 16S rRNA gene was performed using dual-barcoded universal primers 319F and 806R as previously described ^53^. High-throughput sequencing of the amplicons was performed on an Illumina MiSeq platform using the 300 bp paired-end protocol. Metagenomic sequencing libraries were constructed from the same DNA using Illumina Nextera XT kit according to the manufacturer recommendations.

Total RNA was extracted from 250 μl of stool homogenized in RNALater. Briefly, lysis was performed by bead beating using the FastPrep lysing matrix B protocol (MP Biomedicals), followed with two rounds of protein cleanup using phenol-chloroform in 5PRIME heavy phase lock tubes (QuantaBio) and precipitation of total nucleic acids using isopropanol. Genomic DNA was removed using TURBO DNase (Ambion). Ribosomal RNAs were depleted using the Gram-negative and Human/mouse/rat Ribo-Zero rRNA Removal kits (Epicentre Technologies). The resulting RNA was used for library construction using Illumina TruSeq stranded mRNA library preparation kit according to the manufacturer’s recommendations. Quantification of the constructed RNA libraries was performed on an Agilent Bioanalyzer using the DNA 1000 Nano kit. Both metagenome and metatranscriptome samples were sequenced on an Illumina HiSeq 4000 instrument at the Genomics Resource Center (GRC), Institute for Genome Sciences, University of Maryland School of Medicine using the 150 bp paired-end protocol.

### Bioinformatics analysis of intestinal microbiota

Sequencing read quality assessment was performed using strict criteria to ensure high quality and complete sequences of the amplified the V3-V4 regions of the 16S rRNA gene, according to the procedures, programs and citations, and parameters described previously ^53^. Briefly, a sequence read was trimmed at the beginning of a 4 bp sliding window if the average quality score was less than Q15. The sequence read was then assessed for length and retained if it was at least 75% of its original length. The paired-end reads were assembled to take advantage of the ~90bp overlapping region. These sequences were further de-multiplexed the sequence reads by individual samples. Additional quality filtering was applied that removed sequences with more than one mismatch in the barcode sequence tag or with ambiguous nucleotide. Taxonomic assignments were performed on each sequence using the Ribosomal Database Project trained on the Greengene database (Aug 2013 version), using 0.8 confidence values as cutoff. Clustering taxonomic profiles was performed as previously described ^52^. The number of clusters was validated using gap statistics implemented in the *cluster* package in R ^54^ by calculating the goodness of clustering measure. Within-sample diversity was estimated using both observed OTUs to measure community richness and Shannon diversity index. Linear discriminant analysis (LDA) effect size (LEfSe) analysis^55^ was used to identify fecal phylotypes that could explain the differences between infants with low or high La/Rh ratio on different sampling days. For LEfSe, the alpha value for the non-parametric factorial Kruskal-Wallis (KW) sum-rank test was set at 0.05 and the threshold for the logarithmic LDA model ^56^ score for discriminative features was set at 2.0. An all-against-all BLAST search in multi-class analysis was performed.

Balance tree analysis was applied as implemented in Gneiss, and trees were generated using Ward hierarchical clustering of abundance profiles. Balance was computed as the isometric log ratio of mean abundances at each bifurcating node in the tree, to characterize the “flow” of changes in the abundance of a group of correlated bacteria in a microbial community ^20^. Multivariate response linear regression on the calculated balances was performed, and multiple factors were included as covariates, including antibiotics use, maternal antibiotics use, delivery mode, preterm premature rupture of membranes (PPROM), feeding pattern and source, intestinal permeability, birthweight, gender, ethnicity, GA and PMA. Leave-one-variable-out approach was used to calculate the change in R square to evaluate the effect of a single covariate on the community. Ten-fold cross validation was performed to mitigate the common overfitting issues in statistical modelling.

### Statistical Analysis

An adaptive spline logistic regression model implemented in spmrf R package ^57^ was adapted to determine the associations between intestinal permeability and relative abundance of bacterial phylotypes. This model is a locally adaptive nonparametric fitting method that operates within a Bayesian framework, which uses shrinkage prior Markov random fields to induce sparsity and provides a combination of local adaptation and global control ^57^. The analysis was performed on the phylotypes present in at least 15% of all samples, and the effect size was defined as the difference between the extreme values of the probability of intestinal permeability index. Given that there were multiple samples collected from each subject, this model takes into consideration of the dependencies among samples within a subject. Bayesian goodness-of-fit p-value implemented in R package rstan ^58^ was used to access the significance of the association between phylotypes and metadata including antibiotics use, maternal antibiotics use, delivery mode, PPROM, feeding pattern, intestinal permeability, birthweight, gender, ethnicity, gestational age (GA), and postmenstrual age (PMA). R code implementation of the model is provided in **Supplemental File 1**. We further adapted random forest supervised machine learning scheme implemented in R package randomForest ^59^ to test the predictability of the phylotypes of microbial community on intestinal permeability. The top 15 phylotypes relative abundance with highest mean decrease gini index importance measure, were fitted to a random effect logistic regression model of intestinal permeability that was defined as a dichotomous variable high (La/Rh >0.05) or low (La/Rh <=0.05). The relative abundances of phylotypes were centered to the mean and scaled by standard deviation to apply to the model to normalize relative abundances. R code implementation of the model is provided in **Supplemental File 2**.

### Intestinal microbiome analyses

Metagenomic and metatranscriptomic sequence data were pre-processed using the following steps: 1) human sequence reads and rRNA LSU/SSU reads were removed using BMTagger v3.101 ^60^ using a standard human genome reference (GRCh37.p5) ^61^; 2) rRNA sequence reads were removed *in silico* by aligning all reads using Bowtie v1 ^62^ to the SILVA PARC ribosomal-subunit sequence database ^63^. Sequence read pairs were removed even if only one of the reads matched to the human genome reference or to rRNA; 3) the Illumina adapter was trimmed using Trimmomatic ^64^; 4) sequence reads with average quality greater than Q15 over a sliding window of 4 bp were trimmed before the window, assessed for length and removed if less than 75% of the original length; and 5) no ambiguous base pairs were allowed. The taxonomic composition of the microbiomes was established using MetaPhlAn version 2 ^65^. Normalization using Witten-Bell smoothing was performed since metatranscriptomes are a random sampling of all expressed genes and transcripts can be identified that correspond to genes not represented in the metagenome, particularly for low abundance species that were metabolically active ^66^. The relative expression of a gene in a sample was calculated by normalizing the smoothed value of the expression level in the metatranscriptome by the smoothed value of the corresponding gene abundance in the metagenome, as suggested previously ^66,67^. Correlation plots were generated using R *corrplot* package^68^. Genotypic variation of *Escherichia coli* was performed through reconstructing MLST loci-sequences from metagenomes using metaMLST program ^21^. The resulting STs were visualized to show related genotypes of *E. coli* strains on a minimum spanning tree computed by a goeBURST algorithm ^69^ implemented in PHYLOViZ ^70^.

## Conclusion

At birth there is low abundance of *Clostridiales* in preterm infants with progressive, significant increase in abundance in the group with rapid progression toward intestinal barrier maturation, but remained low in those with persistent high IP over the first two weeks of life. We further identified neonatal factors previously identified to promote intestinal barrier maturation, including early exclusive breastmilk feeding and shorter duration antibiotic exposure, favor the early colonization of the gut microbiota by members of the *Clostridiales*, which altogether are associated with improved intestinal barrier function in preterm infants. This highlights the importance of factors such as clinical administration of nutritional supplement and limiting exposure to antibiotic in the high-risk preterm population. Our study suggests rationally selected and formulated *Clostridiales* species could constitute a promising LBP candidate for the prevention of NEC, especially when combined with already available strains of *Bifidobacterium* and *Lactobacillus*. The rationale for this intervention is supported by our correlative finding between increased *Clostridiales* abundance and intestinal barrier maturation of preterm neonates at-risk for NEC development. Identification of specific strains of *Clostridiales,* their functions in mediating intestinal barrier maturation, LBP formulation and manufacturing, dosing, safety and efficacy evaluation will be needed to support their application as oral supplementation to promote intestinal barrier maturation and overall health of preterm neonates. Early prediction and prevention of NEC will ultimately improve overall infant survival rates.

## Data Availability

All 16S rRNA sequence data were deposited in SRA SUB3616368 under BioProject PRJNA432222 (release upon acceptance).

## Contributions

B.M., A.W., A.F., J.R., and R.V. designed the research. B.M., E.M., H.Y., and M.H. performed the research. B.M. and P.G. analyzed the data. B.M., E.M., J.R., and R.V. wrote the paper.

## Acknowledgements

This study was funded by The Gerber Foundation and NCCIH (National Center for Complementary and Integrative Health, AT006945).

This work is dedicated to the memory of our colleague Bushra Saleem, M.B.B.S., who contributed to the design and conduct of the study.

The authors thank Dr. Emmanuel Mongodin and Dr. Lauren Hittle, PhD at the Institute for Genome Sciences - University of Maryland School of Medicine for their helpful assistance in total RNA extraction.

## Competing interest statement

The authors declare no competing financial and non-financial interests.

## References

1 Lee, S. H. Intestinal permeability regulation by tight junction: implication on inflammatory bowel diseases. Intest Res 13, 11–18, doi:10.5217/ir.2015.13.1.11 (2015).

2 Fasano, A. Physiological, pathological, and therapeutic implications of zonulin-mediated intestinal barrier modulation: living life on the edge of the wall. Am J Pathol 173, 1243–1252, doi:10.2353/ajpath.2008.080192 (2008).

3 Neish, A. S. Microbes in gastrointestinal health and disease. Gastroenterology 136, 65–80, doi:10.1053/j.gastro.2008.10.080 (2009).

4 Sharon, I. et al. Time series community genomics analysis reveals rapid shifts in bacterial species, strains, and phage during infant gut colonization. Genome Res 23, 111–120, doi:10.1101/gr.142315.112 (2013).

5 Belkaid, Y. & Hand, T. W. Role of the microbiota in immunity and inflammation. Cell 157, 121–141, doi:10.1016/j.cell.2014.03.011 (2014).

6 Arrieta, M. C., Stiemsma, L. T., Amenyogbe, N., Brown, E. M. & Finlay, B. The intestinal microbiome in early life: health and disease. Front Immunol 5, 427, doi:10.3389/fimmu.2014.00427 (2014).

7 Madan, J. C., Farzan, S. F., Hibberd, P. L. & Karagas, M. R. Normal neonatal microbiome variation in relation to environmental factors, infection and allergy. Curr Opin Pediatr 24, 753–759, doi:10.1097/MOP.0b013e32835a1ac8 (2012).

8 Arrieta, M. C. et al. Early infancy microbial and metabolic alterations affect risk of childhood asthma. Sci Transl Med 7, 307ra152, doi:10.1126/scitranslmed.aab2271 (2015).

9 Vatanen, T. et al. Variation in Microbiome LPS Immunogenicity Contributes to Autoimmunity in Humans. Cell 165, 842–853, doi:10.1016/j.cell.2016.04.007 (2016).

10 Cenit, M. C., Olivares, M., Codoner-Franch, P. & Sanz, Y. Intestinal Microbiota and Celiac Disease: Cause, Consequence or Co-Evolution? Nutrients 7, 6900–6923, doi:10.3390/nu7085314 (2015).

11 Gevers, D. et al. The treatment-naive microbiome in new-onset Crohn’s disease. Cell Host Microbe 15, 382–392, doi:10.1016/j.chom.2014.02.005 (2014).

12 Cho, I. et al. Antibiotics in early life alter the murine colonic microbiome and adiposity. Nature 488, 621–626, doi:10.1038/nature11400 (2012).

13 Guner, Y. S. et al. State-based analysis of necrotizing enterocolitis outcomes. J Surg Res 157, 21–29, doi:10.1016/j.jss.2008.11.008 (2009).

14 Fitzgibbons, S. C. et al. Mortality of necrotizing enterocolitis expressed by birth weight categories. J Pediatr Surg 44, 1072–1075; discussion 1075-1076, doi:S0022-3468(09)00160-2 [pii] 10.1016/j.jpedsurg.2009.02.013 (2009).

15 Fox, T. P. & Godavitarne, C. What really causes necrotising enterocolitis? ISRN Gastroenterol 2012, 628317, doi:10.5402/2012/628317 (2012).

16 Saleem, B. et al. Intestinal Barrier Maturation in Very Low Birthweight Infants: Relationship to Feeding and Antibiotic Exposure. J Pediatr 183, 31–36 e31, doi:10.1016/j.jpeds.2017.01.013 (2017).

17 Mshvildadze, M., Neu, J. & Mai, V. Intestinal microbiota development in the premature neonate: establishment of a lasting commensal relationship? Nutrition reviews 66, 658–663, doi:10.1111/j.1753-4887.2008.00119.x (2008).

18 Mackie, R. I., Sghir, A. & Gaskins, H. R. Developmental microbial ecology of the neonatal gastrointestinal tract. Am J Clin Nutr 69, 1035S–1045S (1999).

19 Unger, S., Stintzi, A., Shah, P., Mack, D. & O’Connor, D. L. Gut microbiota of the very-low-birth-weight infant. Pediatr Res 77, 205–213, doi:10.1038/pr.2014.162 (2015).

20 Morton, J. T. et al. Balance Trees Reveal Microbial Niche Differentiation. mSystems 2, doi:10.1128/mSystems.00162-16 (2017).

21 Zolfo, M., Tett, A., Jousson, O., Donati, C. & Segata, N. MetaMLST: multi-locus strain-level bacterial typing from metagenomic samples. Nucleic Acids Res 45, e7, doi:10.1093/nar/gkw837 (2017).

22 Guner, Y. S., Malhotra, A., Ford, H. R., Stein, J. E. & Kelly, L. K. Association of Escherichia coli O157:H7 with necrotizing enterocolitis in a full-term infant. Pediatr Surg Int 25, 459–463, doi:10.1007/s00383-009-2365-3 (2009).

23 Ward, D. V. et al. Metagenomic Sequencing with Strain-Level Resolution Implicates Uropathogenic E. coli in Necrotizing Enterocolitis and Mortality in Preterm Infants. Cell Rep 14, 2912–2924, doi:10.1016/j.celrep.2016.03.015 (2016).

24 Palmer, C., Bik, E. M., Digiulio, D. B., Relman, D. A. & Brown, P. O. Development of the Human Infant Intestinal Microbiota. PLoS Biol 5, e177 (2007).

25 Neu, J. & Walker, W. A. Necrotizing enterocolitis. N Engl J Med 364, 255–264, doi:10.1056/NEJMra1005408 (2011).

26 Mai, V. et al. Fecal microbiota in premature infants prior to necrotizing enterocolitis. PLoS One 6, e20647, doi:10.1371/journal.pone.0020647 (2011).

27 Taylor, S. N., Basile, L. A., Ebeling, M. & Wagner, C. L. Intestinal permeability in preterm infants by feeding type: mother’s milk versus formula. Breastfeed Med 4, 11–15, doi:10.1089/bfm.2008.0114 (2009).

28 Bergmann, K. R. et al. Bifidobacteria stabilize claudins at tight junctions and prevent intestinal barrier dysfunction in mouse necrotizing enterocolitis. Am J Pathol 182, 1595–1606, doi:10.1016/j.ajpath.2013.01.013 (2013).

29 Nanthakumar, N. et al. The mechanism of excessive intestinal inflammation in necrotizing enterocolitis: an immature innate immune response. PLoS One 6, e17776, doi:10.1371/journal.pone.0017776 (2011).

30 van Elburg, R. M., Fetter, W. P., Bunkers, C. M. & Heymans, H. S. Intestinal permeability in relation to birth weight and gestational and postnatal age. Arch Dis Child Fetal Neonatal Ed 88, F52–55 (2003).

31 Carlisle, E. M. & Morowitz, M. J. The intestinal microbiome and necrotizing enterocolitis. Curr Opin Pediatr 25, 382–387, doi:10.1097/MOP.0b013e3283600e91 (2013).

32 La Rosa, P. S. et al. Patterned progression of bacterial populations in the premature infant gut. Proc Natl Acad Sci U S A 111, 12522–12527, doi:10.1073/pnas.1409497111 (2014).

33 Barron, L. K. et al. Independence of gut bacterial content and neonatal necrotizing enterocolitis severity. J Pediatr Surg 52, 993–998, doi:10.1016/j.jpedsurg.2017.03.029 (2017).

34 Penders, J. et al. Factors influencing the composition of the intestinal microbiota in early infancy. Pediatrics 118, 511–521, doi:10.1542/peds.2005-2824 (2006).

35 Colaizy, T. T. et al. Impact of Optimized Breastfeeding on the Costs of Necrotizing Enterocolitis in Extremely Low Birthweight Infants. J Pediatr 175, 100–105 e102, doi:10.1016/j.jpeds.2016.03.040 (2016).

36 Narushima, S. et al. Characterization of the 17 strains of regulatory T cell-inducing human-derived Clostridia. Gut microbes 5, 333–339, doi:10.4161/gmic.28572 (2014).

37 Blakely, M. L. et al. Postoperative outcomes of extremely low birth-weight infants with necrotizing enterocolitis or isolated intestinal perforation: a prospective cohort study by the NICHD Neonatal Research Network. Ann Surg 241, 984–989; discussion 989-994 (2005).

38 Stratiki, Z. et al. The effect of a bifidobacter supplemented bovine milk on intestinal permeability of preterm infants. Early Hum Dev 83, 575–579, doi:10.1016/j.earlhumdev.2006.12.002 (2007).

39 Deshpande, G., Rao, S., Patole, S. & Bulsara, M. Updated meta-analysis of probiotics for preventing necrotizing enterocolitis in preterm neonates. Pediatrics 125, 921–930, doi:peds.2009-1301 [pii] 10.1542/peds.2009-1301 (2010).

40 Stefka, A. T. et al. Commensal bacteria protect against food allergen sensitization. Proc Natl Acad Sci U S A 111, 13145–13150, doi:10.1073/pnas.1412008111 (2014).

41 Rajilic-Stojanovic, M., Smidt, H. & de Vos, W. M. Diversity of the human gastrointestinal tract microbiota revisited. Environ Microbiol 9, 2125–2136, doi:10.1111/j.1462- 2920.2007.01369.x (2007).

42 Walker, A. W. et al. Dominant and diet-responsive groups of bacteria within the human colonic microbiota. Isme J 5, 220–230, doi:10.1038/ismej.2010.118 (2011).

43 Khanna, S. et al. A Novel Microbiome Therapeutic Increases Gut Microbial Diversity and Prevents Recurrent Clostridium difficile Infection. J Infect Dis 214, 173–181, doi:10.1093/infdis/jiv766 (2016).

44 Smith, P. M. et al. The microbial metabolites, short-chain fatty acids, regulate colonic Treg cell homeostasis. Science 341, 569–573, doi:10.1126/science.1241165 (2013).

45 Atarashi, K. et al. Induction of colonic regulatory T cells by indigenous Clostridium species. Science 331, 337–341, doi:10.1126/science.1198469 (2011).

46 Atarashi, K. et al. Treg induction by a rationally selected mixture of Clostridia strains from the human microbiota. Nature 500, 232–236, doi:10.1038/nature12331 (2013).

47 McMurtry, V. E. et al. Bacterial diversity and Clostridia abundance decrease with increasing severity of necrotizing enterocolitis. Microbiome 3, 11, doi:10.1186/s40168-015-0075-8 (2015).

48 Kim, Y. G. et al. Neonatal acquisition of Clostridia species protects against colonization by bacterial pathogens. Science 356, 315–319, doi:10.1126/science.aag2029 (2017).

49 Jovel, J. et al. Characterization of the Gut Microbiome Using 16S or Shotgun Metagenomics. Frontiers in microbiology 7, 459, doi:10.3389/fmicb.2016.00459 (2016).

50 Grier, A. et al. Impact of prematurity and nutrition on the developing gut microbiome and preterm infant growth. Microbiome 5, 158, doi:10.1186/s40168-017-0377-0 (2017).

51 Sellitto, M. et al. Proof of concept of microbiome-metabolome analysis and delayed gluten exposure on celiac disease autoimmunity in genetically at-risk infants. PLoS One 7, e33387, doi:10.1371/journal.pone.0033387 (2012).

52 Ravel, J. et al. Vaginal microbiome of reproductive-age women. Proc. Natl. Acad. Sci. USA 108 Suppl 1, 4680–4687, doi:10.1073/pnas.1002611107 (2011).

53 Fadrosh, D. W. et al. An improved dual-indexing approach for multiplexed 16S rRNA gene sequencing on the Illumina MiSeq platform. Microbiome 2, 6, doi:10.1186/2049-2618-2-6 (2014).

54 Maechler, M. cluster: "Finding Groups in Data": Cluster Analysis Extended Rousseeuw et al., 2016).

55 Segata, N. et al. Metagenomic biomarker discovery and explanation. Genome Biol 12, R60, doi:10.1186/gb-2011-12-6-r60 (2011).

56 Fisher, R. A. The use of multiple measurements in taxonomic problems. Ann Eugenics 7 (1936).

57 Faulkner, J. R., Minin, V. Locally adaptive smoothing with Markov random fields and shrinkage priors. Bayesian Analysis 13, 225–252 (2018).

58 Team, S. D. RStan: the R interface to Stan. R package version 2.17.3. (2018).

59 Liaw, A. & Wiener, M. Classification and Regression by randomForest.. R News 2, 18–22 (2002).

60 Rotmistrovsky, K. & Agarwala, R. BMTagger: Best Match Tagger for removing human reads from metagenomics datasets (NCBI/NLM, National Institutes of Health, 2011).

61 Church, D. M. et al. Modernizing reference genome assemblies. PLoS Biol 9, e1001091, doi:10.1371/journal.pbio.1001091 (2011).

62 Langmead, B., Trapnell, C., Pop, M. & Salzberg, S. L. Ultrafast and memory-efficient alignment of short DNA sequences to the human genome. Genome Biol 10, R25, doi:10.1186/gb-2009-10-3-r25 (2009).

63 Quast, C. et al. The SILVA ribosomal RNA gene database project: improved data processing and web-based tools. Nucleic Acids Res 41, D590–596, doi:10.1093/nar/gks1219 (2013).

64 Bolger, A. M., Lohse, M. & Usadel, B. Trimmomatic: a flexible trimmer for Illumina sequence data. Bioinformatics 30, 2114–2120, doi:10.1093/bioinformatics/btu170 (2014).

65 Segata, N. et al. Metagenomic microbial community profiling using unique clade-specific marker genes. Nat Methods 9, 811–814, doi:10.1038/nmeth.2066 (2012).

66 Franzosa, E. A. et al. Relating the metatranscriptome and metagenome of the human gut. Proc Natl Acad Sci U S A 111, E2329–2338, doi:10.1073/pnas.1319284111 (2014).

67 Franzosa, E. A. et al. Sequencing and beyond: integrating molecular ‘omics’ for microbial community profiling. Nat Rev Microbiol 13, 360–372, doi:10.1038/nrmicro3451 (2015).

68 Wei, T. & Simko, V. R package "corrplot": Visualization of a Correlation Matrix. (2017).

69 Francisco, A. P., Bugalho, M., Ramirez, M. & Carrico, J. A. Global optimal eBURST analysis of multilocus typing data using a graphic matroid approach. BMC Bioinformatics 10, 152, doi:10.1186/1471-2105-10-152 (2009).

70 Nascimento, M. et al. PHYLOViZ 2.0: providing scalable data integration and visualization for multiple phylogenetic inference methods. Bioinformatics 33, 128–129, doi:10.1093/bioinformatics/btw582 (2017).

71 Oksanen, J. et al. vegan: Community Ecology Package. R package version 2.0-2. (2011).

